# A 340/380 nm light emitting diode illuminator for Fura-2 AM ratiometric Ca^2+^ imaging of live cells with better than 5 nM precision

**DOI:** 10.1101/138495

**Authors:** Peter W. Tinning, Aimee J. P.M. Franssen, Shehla U. Hridi, Trevor J. Bushell, Gail McConnell

## Abstract

We report the first demonstration of a fast wavelength-switchable 340/380 nm light emitting diode (LED) illuminator for Fura-2 ratiometric Ca^2+^ imaging of live cells. The LEDs closely match the excitation peaks of bound and free Fura-2 and enables the precise detection of cytosolic Ca^2+^ concentrations, which is only limited by the Ca^2+^ response of Fura-2. Using this illuminator, we have shown that Fura-2 acetoxymethyl ester (AM) concentrations as low as 250 nM can be used to detect induced Ca^2+^ events in tsA-201 cells and while utilizing the 150 μs switching speeds available, it was possible to image spontaneous Ca^2+^ transients in hippocampal neurons at a rate of 24.39 Hz that were blunted or absent at typical 0.5 Hz acquisition rates. Overall, the sensitivity and acquisition speeds available using this LED illuminator significantly improves the temporal resolution that can be obtained in comparison to current systems and supports optical imaging of fast Ca^2+^ events using Fura-2.

## Introduction

Calcium (Ca^2+^) plays a varied and integral role in mediating and controlling many biological processes including the regulation of muscle contractions [1], triggering insulin release from pancreatic cells [2], and the release of neurotransmitters in neurons [3]. Increases in cytosolic Ca^2+^ levels can originate from a number of sources including release from internal stores triggered by activation of G-protein coupled receptors by both endogenous and exogenous stimuli [4], or from external sources via influx through voltage-gated Ca^2+^ channels [5]. Hence, the measurement of Ca^2+^ dynamics has been utilized extensively in biological research as it can reveal how specimens respond to different stimuli and how Ca^2+^ signaling is altered in disease states.

Whilst electrophysiological studies are still the gold standard for measuring electrical activity within and between excitable cells due to their high temporal resolution [6], the spatial resolution is low and the nature of the technique leads to low throughput data production. In contrast, the development of Ca^2+^ specific fluorescent indicators allows for high throughput data acquisition with good spatial resolution [6]–[8], which has allowed intracellular Ca^2+^ dynamics to be investigated non-invasively in multiple cells simultaneously using widefield epifluorescence microscopy.

Fluorescent Ca^2+^ indicators typically fall into two different categories, namely single excitation wavelength indicators, including Fluo-4 and Fluo-3, or dual-wavelength dyes (emission or excitation) such as Fura-2 or lndo-1 [9], [10]. Single wavelength indicators have a high quantum yield, allow for a simple excitation and detection setup. The Ca^2+^ concentration changes are then identifiable through an emission intensity change [11]. However, these indicators are unable to provide quantitative Ca^2+^ data since the emission intensities may be influenced by dye concentration or photobleaching during imaging [9], [12],

Dual wavelength or ratiometric Ca^2+^ indicators have either excitation or emission wavelengths that shift in response to concentration changes in cytosolic Ca^2+^ [13], and as such require an imaging setup that ensures that the free and bound Ca^2+^ wavelengths are recorded separately. The quantitative cytosolic Ca^2+^concentrations are obtained by taking a ratio of the Ca^2+^ free and bound wavelengths, with these ratios being unaffected by the light intensities or the dye concentration within the cytosol [11], [14], [15]. Whilst quantitative data can be acquired, dual wavelength indicators typically have a smaller dynamic range than single wavelength dyes [9]. Fura-2 is a ratiometric fluorescent Ca^2+^ indicator that was developed as an improved alternative to the Ca^2+^ indicator, Quin2 [16]. Fura-2 also holds advantages over another ratiometric dye, lndo-1, as it has a larger dynamic range between Ca^2+^ bound and free states [9], and is more resistant to photobleaching [11]. When cytosolic free Ca^2+^ binds to Fura-2, the peak excitation wavelength changes from 380 nm to 340 nm whilst the peak emission around 510 nm remains unchanged [16]. By sequential excitation of Fura-2 at 340 nm and 380 nm and taking a ratio of the emission signals for each excitation wavelength, these ratios can be calibrated to a measurement of the corresponding cytosolic Ca^2+^ concentration by measuring the ratio of the fluorescence emission signal in the presence of known free Ca^2+^ concentrations.

Historically, the most commonly used light source for widefield Fura-2 excitation has been an arc lamp with a monochromator [11]. Users of these systems have had to sacrifice precise and immediate control over light intensity without the use of neutral density filters and are limited to wavelength switching speeds on a millisecond timescale. In addition, arc lamp light sources exhibit inherent amplitude instability on the order of 5% [17], which reduces the accuracy of measurement and as a result small changes in Ca^2+^ may go undetected [18]– [20].

Previous investigations have used two-photon microscopy to conduct Fura-2 Ca^2+^ imaging [21], [22]. By using this technique, it is possible to reduce photo-bleaching rates in the out of focus planes [23] and image deeper into bulk specimens [24]. However, point-scanning two-photon excitation is slow [25], [26], as is the wavelength scanning of the laser [27]. This combination does not readily support detection and measurement of fast changes in Ca^2+^ concentration that are possible with a widefield microscope and a fast camera. More recently, widefield two photon microscopy has been shown to image hippocampal neurons loaded with Fluo-4 [28], but this work does not support quantitative measurement of Ca^2+^concentration.

More recently, light emitting diodes (LEDs) have been utilized to excite Fura-2. This type of illuminator can support high stability switching on microsecond timescales and offer precise output intensity control by simply changing the LED drive current. Until recently, commercial LED systems have only offered LED combinations of 350/380 nm or 360/380 nm, which do not precisely match the excitation wavelengths required or only allow excitation at the isosbestic point [9]. We have taken advantage of a new shorter wavelength, high-brightness LED at 340 nm to develop a 340/380 nm switchable LED illuminator. We demonstrate its application in microscopy by performing Fura-2 ratiometric Ca^2+^ imaging in both an immortalized cell line and in primary cultured neurons exhibiting pharmacologically-induced and synaptically-driven Ca^2+^ responses.

## Materials and methods

### Characterization of 340/380 nm LED system

The peak output spectra for the 340 nm (pE-100-340, CoolLED, Andover, UK) and 380 nm LEDs (pE-100-380, CoolLED, Andover, UK) were measured using a spectrometer (USB2000+UV-VIS-ES, OceanOptics, Florida, USA). The peak wavelength and full width at half maximum (FWHM) for each LED were found to be 342.2 ± 1.5 (FWHM: 9.3 ± 1.5) nm and 383.3 ± 1.5 (FWHM: 8.3 ± 1.5) nm. Plots of these spectra can be seen in Figure 1.

**Figure 1.**
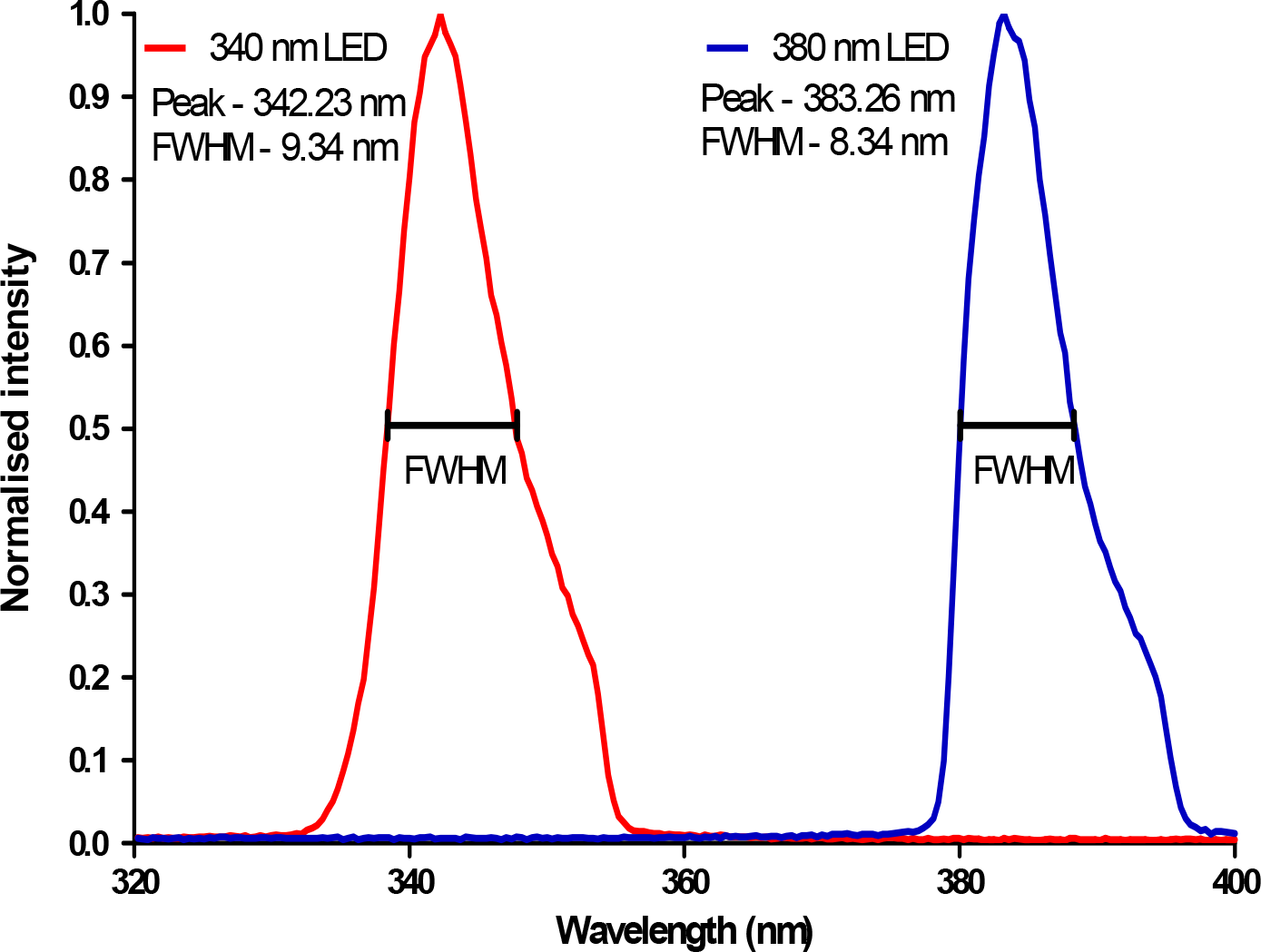
Output spectra of 340/380 nm LEDs obtained at a drive current of 1.52 ± 0.16 A

Power measurements were recorded using a power meter (Fieldmax II, Coherent, California, USA) with a thermal head (PM10, Coherent, California, USA). The average power was taken from three separate measurements with an integration time of 3 seconds. Measurements were taken at the specimen plane under an Olympus 20X/0.5 water dipping objective lens at drive currents up to 1.52 ± 0.16 A for each LED.

The 340 nm LED demonstrated a linear increase in optical power of approximately 6.6 mW/A up to 0.59 ± 0.07 A. Above this current the 340 nm LED exhibited rollover, a phenomenon where with an increase in drive current the optical power begins to plateau or even decrease. The 380 nm LED showed a linear increase in optical power of approximately 14.7 mW/A increase in current up to 1.52 ± 0.16 A. The optical power at different drive currents are shown in Figure 2.

**Figure 2.**
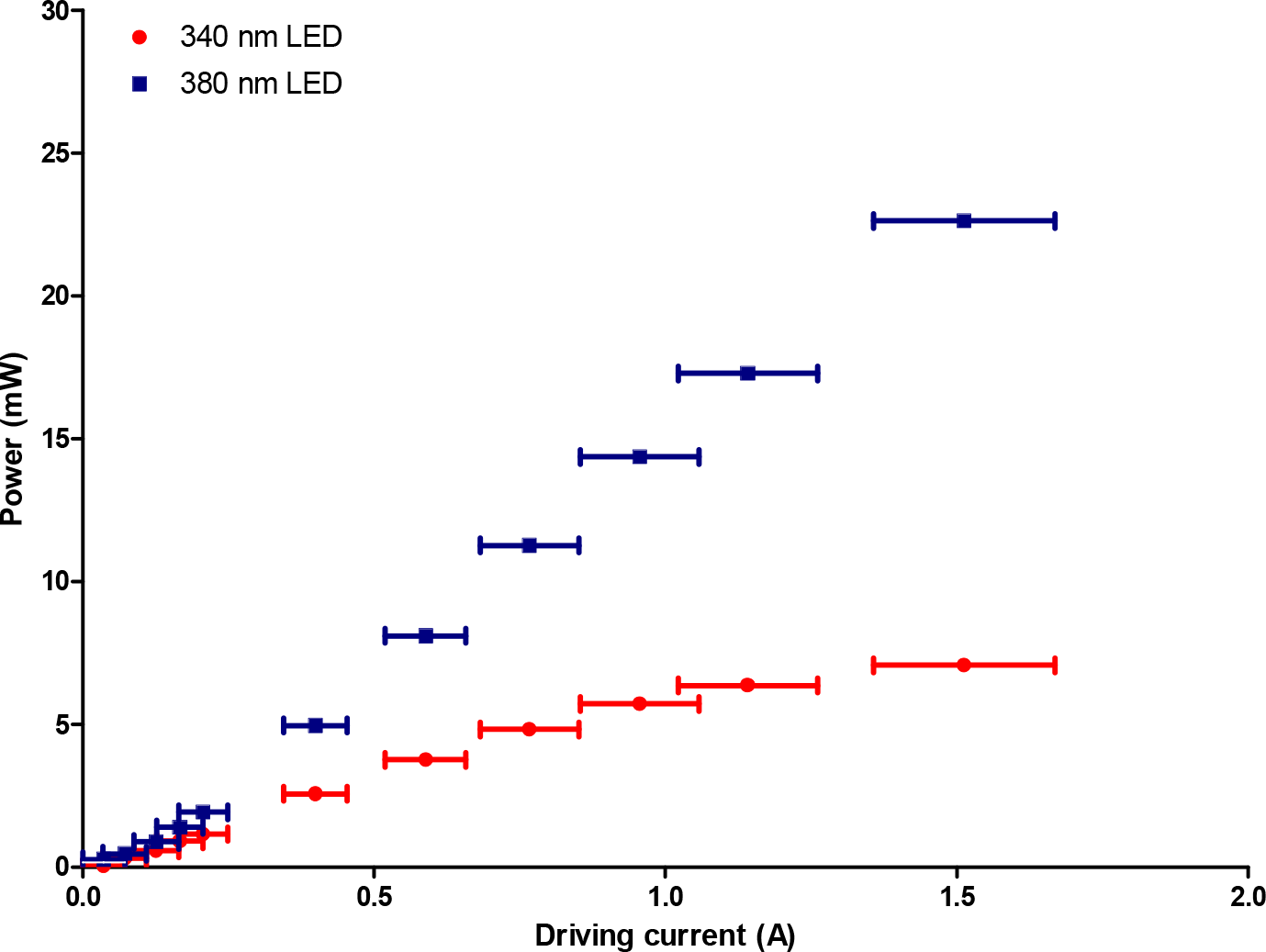
Optical powers measured at the specimen plane of an Olympus BX50 microscope under a 20X water dipping lens at different LED drive currents.

The 340 nm LED demonstrated a linear increase in optical power of approximately 6.6 mW/A up to 0.59 ± 0.07 A. Above this current the 340 nm LED exhibited rollover, a phenomenon where with an increase in drive current the optical power begins to plateau or even decrease. The 380 nm LED showed a linear increase in optical power of approximately 14.7 mW/A increase in current up to 1.52 ± 0.16 A. The optical power at different drive currents are shown in Figure 2.

The 340 nm and 380 nm LEDs had an average power at the specimen plane of 3.76 ± 0.02 mW and 8.10 ± 0.03 mW respectively when driven at a current of 0.59 ± 0.07 A, which was the largest driving current measured before the 340 nm LED begins to rollover. For the experiments presented, the 340 nm LED was used at an optical power at the microscope specimen plane of 1.32 ± 0.01 mW and the 380 nm LED was kept between 1.40 ± 0.02 mW and 3.08 ± 0.01 mW. The Olympus 20X/0.5 water dipping objective lens gives a field size of 0.95 mm^2^ so using these LED output power values we measure intensities at the specimen plane of 1.39 ± 0.01 mW/mm^2^for the 340 nm LED and between 1.47 ± 0.02 mW/mm^2^ and 3.24 ± 0.01 mW/mm^2^for the 380 nm LED.

### tsA-201 cell culture

tsA-201 cells, a modified HEK-293 cell line, were maintained in 5% CO2, in a humidified incubator at 37 °C, in growth media containing Minimum Essential Media (MEM) (with Earle’s, without L-glutamine), 10% foetal calf serum (Biosera, Sussex, UK), 1% non-essential amino acids (Gibco, Paisley, UK), 1% penicillin (10,000 U/ml) and streptomycin (10 mg/ml) (Sigma, Dorset, UK). When the cells were 90% confluent, they were split and plated on 13mm glass coverslips coated with poly-L-lysine (O.lmg.ml’^1^, Sigma, Dorset, UK).

### Primary mouse hippocampal culture

Mouse hippocampal cultures were prepared as described previously (Gan *et al.*, 2011; Abdul Rahman *et al.*, 2016). Briefly, 1 to 2 day-old C57/BL6J pups were killed by cervical dislocation and decapitated. The hippocampi were removed, triturated and the resulting cells were plated out at a density of 5.5 x 10^5^ cells.mL^-1^ onto 13 mm poly-L-lysine (O.Olmg.ml^1^) coated coverslips. Cultures were incubated in culture medium consisting of Neurobasal-A Medium (Invitrogen, Paisley, UK) supplemented with 2% (v/v) B-27 (Invitrogen, Paisley, UK) and 2 mM L-glutamine and maintained in a humidified atmosphere at 37°C/5% CO_2_ for 10–14 days in vitro (DIV). All animal care and experimental procedures were in accordance with UK Home Office guidelines and approved by the University of Strathclyde Ethics Committee.

### Ca^2+^ imaging

Cultures were washed three times in HEPES-buffered saline (HBS) composed of the following (in mM): NaCl 140, KCl 5, MgCl_2_ 2, CaCl_2_ 2, HEPES 10, D-glucose 10, pH 7.4, 310 ± 2 mOsm. They were then loaded with Fura-2 AM (250 nM - 1 µM; Invitrogen, Paisley, UK) made up in HBS for 45 – 60 minutes at 37 °C, after which they were washed with HBS a further three times prior to imaging. Throughout imaging, the cultures were constantly perfused with HBS at a rate of 3 – 3.5 ml.min ^-1^ with all drug solutions being added via the perfusate. Cultures were placed in a perfusion bath under a 20x/0.5 water dipping objective lens (Olympus UMPlanFl) in an upright widefield epifluorescence microscope (Olympus BX50). The 340 nm and 380 nm LEDs were coupled to the microscope with a beam path combiner (pE-combiner, CoolLED) which contained a >365 dichroic mirror (365dmlp, Chroma), and clean up filters for each LED (Semrock BrightLine 340/26 nm bandpass filter for the 340 nm LED and a Semrock BrightLine 387/11 nm bandpass filter for the 380 nm LED). A schematic diagram for this setup can be seen below (Figure 3).

**Figure 3.**
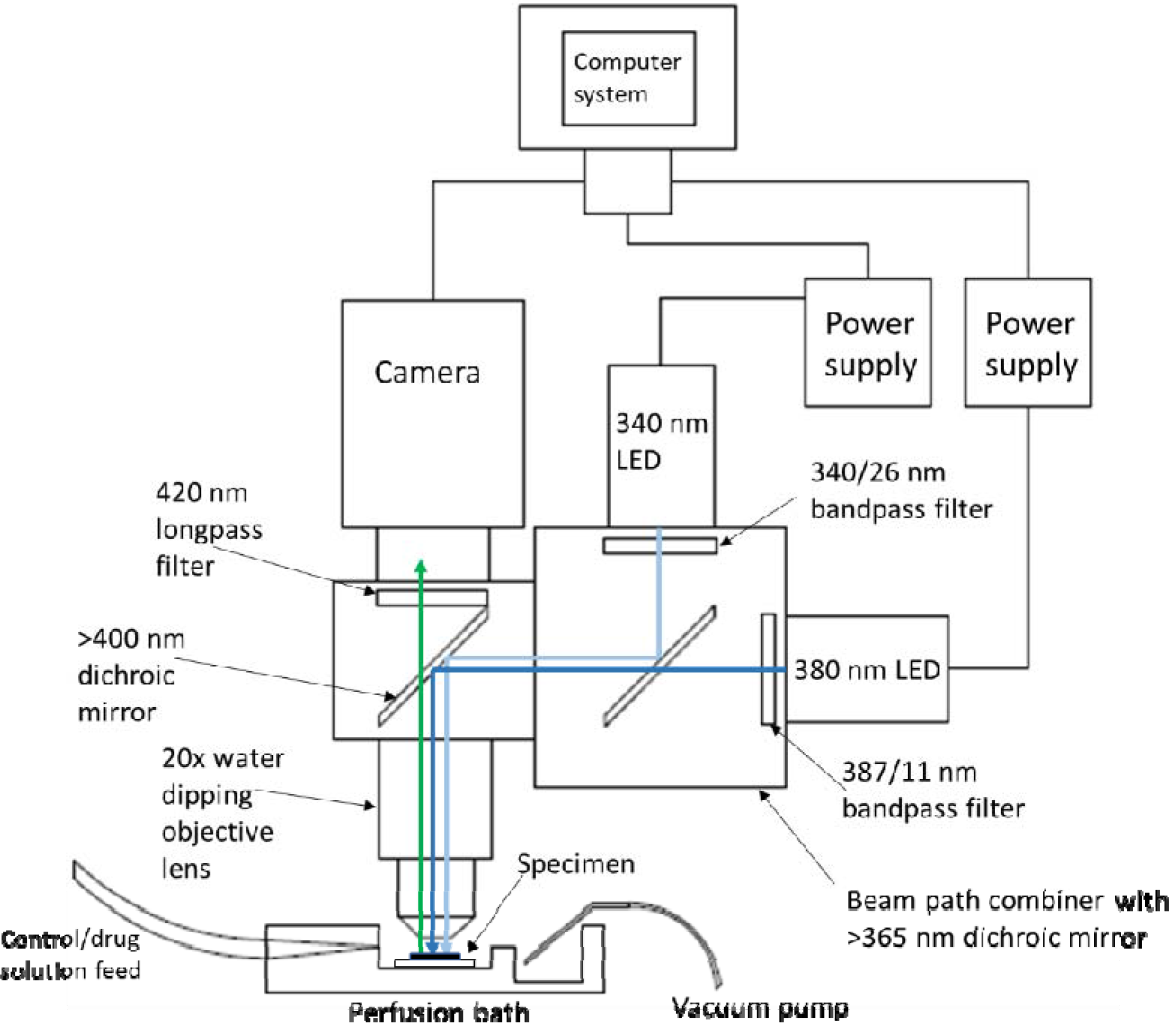
Schematic diagram of experimental imaging setup showing the location of the specimen in relation to the objective lens and where the perfused solution flows over the specimen and gets removed from the bath. The light paths of the 340/330 nm LEDs are also shown to converge through the use of 365 nm dichroic mirror and then illuminate the specimen in the perfusion bath seguentially. The emitted Fura-2 AM fluorescence propagates upwards through the objective lens, >400 nm dichroic mirror (Olympus) and 420 longpass filter (Olympus) to the camera. The camera and power supplies for both LEDs are connected to a computer system for TTL triggering and recording fluorescent signals

During routine imaging, cultures were alternatively illuminated with the 340 nm and 380 nm LEDs, each with an exposure time of 100 ms, and imaged at a rate of 0.5 Hz with emission being detected above 420 nm by a CMOS camera (ORCA-Flash 4.0, Hamamatsu) using a binning of n=2. For video rate image capture, an LED switching speed of 150 µs was used with exposure times reduced to 20.5 ms, facilitating imaging at a rate of 24.39 Hz. All signals were recorded using the WinFluor imaging software[31], which also synchronized and TTL triggered the 340/380 nm illuminator and camera. Results were calculated as changes in fluorescence ratio occurring within the cytosol and expressed as a fold increase above the normalized baseline.

### Data analysis and statistics

An area on each coverslip which was free of cells was selected to determine the background fluorescence level and subtracted from each of the ROIs. The background-corrected emission fluorescence time courses from 340 nm and 380 nm excitation obtained using WinFluor were read into a MATLAB script to determine the average baseline Ca^2+^ level obtained during the initial HBS solution wash. Each ROI was then normalized to the associated calculated baseline. The average peak fold increases of emission ratios above the baseline for drug washes were then calculated using the normalized emission ratios.

To convert the emission ratios from the 340/380 nm excitation into a measurement of the free cytosolic Ca^2+^ concentration we used the following equation[16],

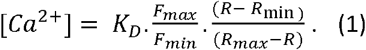

where *K_D_* is the dissociation constant for Fura-2 (224 nM) [16], *R* is the experimental emission ratios and *F_max_, R_max_ F_min_* and *R_min_* are the 380 nm fluorescence emission signals and emission ratio from 340/380 nm excitation at saturating and zero free Ca^2+^ levels, respectively. To determine the experimental values required for equation (1) we recorded the emission ratios obtained from 39 µM and 0 µM free Ca^2+^ solutions obtained using a Fura-2 Ca^2+^ imaging calibration kit (Invitrogen, Paisley, UK). The percentage Ca^2+^ baseline fluctuations were determined by finding the maximum and minimum deviations from the recorded baseline. This deviation was then calculated as the percentage of the average baseline level.

All biological replicates are reported as an ‘n’ number which are equal to the total number of ROIs investigated throughout each experiment taken from at least 4 different cultures. All data are expressed as mean ± standard error of the mean. Data were compared by using either an unpaired student t-test or a one-way ANOVA with Tukey’s comparison when appropriate, with P values < 0.05 considered significant.

## Results

### Fura-2 AM ratiometric Ca^2+^ imaging of induced Ca^2+^ transients in live cell specimens

Pharmacologically-induced increases in intracellular Ca^2+^ were observed in tsA-201 cells (n = 572) and cultured hippocampal neurons (n = 388). The pharmacological stimuli were selected for use as they are known to cause large but slow Ca^2+^ concentration changes in the live cell specimens. The normalized emission ratio fold increases above the resting baseline in the tsA-201 cells were 1.67 ± 0.04 (n = 572) evoked by ATP (5 µM) and 3.08 ± 0.04 (n = 572) by trypsin (100 nM), respectively. In hippocampal neurons, application of glutamate (20 µM) caused a fluorescence fold increase of 4.2 ± 0.1 (n = 388) with potassium chloride (KCI, 20 Mm) application eliciting fold increases of 2.51 ± 0.06 (n = 388).

Using equation (1) each ROI was converted to a measurement of cytosolic Ca^2+^to allow quantitative data to be obtained. From measurement of the cytosolic Ca^2+^ concentrations using Ca^2+^standards, ATP (5 µM) induced cytosolic Ca^2+^ increases of 280.4 ± 7.8 nM (n = 572, P<O.OOOl compared to average baseline of 81.2 ± 5.6 nM, Figure 4A and B,) and trypsin (100 nM) caused a 581.9 ± 10.2 nM increase (n = 572, P<O.OOOl compared to average baseline of 81.6 ± 5.6 nM, Figure 4A and B) in the tsA-201 cells. In hippocampal neurons, glutamate (20 µM) induced Ca^2+^ increases of 645.4 ± 18.2 nM (n = 388, P<O.OOOl compared to average baseline of 92.3 ± 10.1 nM, Figure 4C and D) and KCl (20 mM) elicited increases in Ca^2+^ of 357.6 ± 9.2 nM (n = 388, P<O.OOOl compared to baseline of 92.3 ± 10.1 nM, Figure 4C and D).

**Figure 4.**
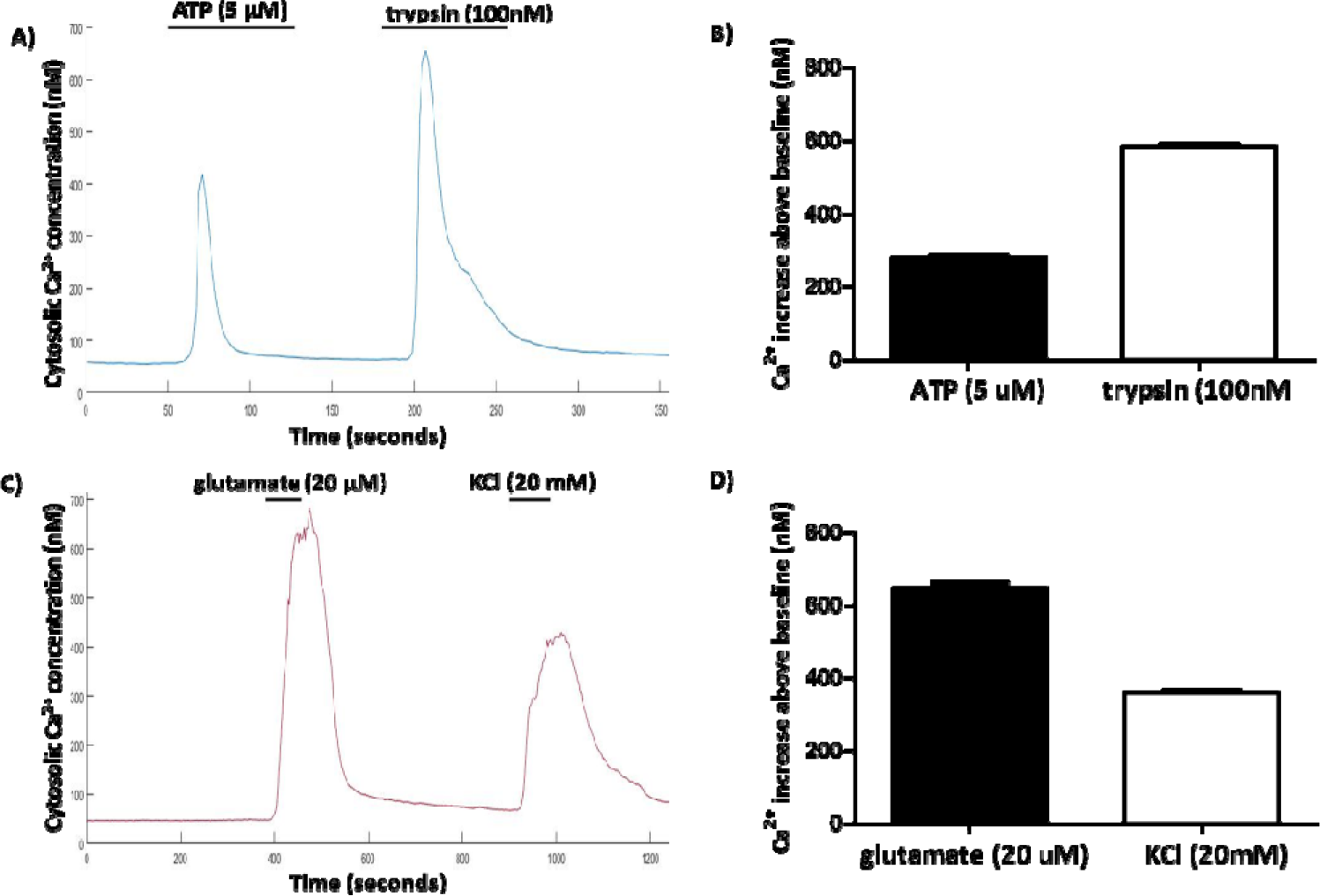
A) Representative trace of Ca2+changes in tsA-201 cells (n = 572) elicited by the application of ATP (5 μM) and trypsin (100 nM). B) Average pharmacologically stimulated Ca2+ increases above the baseline levels in tsA-201 cells. C) Representative trace of Ca2+ increase in hippocampal neurons (n = 388) through the application of glutamate (20 μM) and KCl (20 mM). D) Average Ca2+ increases above resting levels in hippocampal neurons for each stimulus.

### Ca^2+^ baseline fluctuation measurements

To determine the minimum cytosolic Ca^2+^ concentration change that could be accurately monitored from the baseline levels using the 340/380 nm LED illuminator, we analyzed the baseline fluctuations observed in the experiments reported above. By obtaining the maximum and minimum values in each of the baseline measurements for the tsA-201 and hippocampal neuron Ca^2+^ imaging experiments, we measured an average peak to peak fluctuation of 5.9 ± 0.2 % (n = 572) for the tsA-201 cells and 4.2 ± 0.2 % in the hippocampal neurons. Using equation (1), these fluctuations equate to a cytosolic Ca^2+^ concentration fluctuation of 4.8 ± 0.2 nM in the tsA-201 cells and 3.9 ± 0.2 nM in the hippocampal neurons.

**Figure 5.**
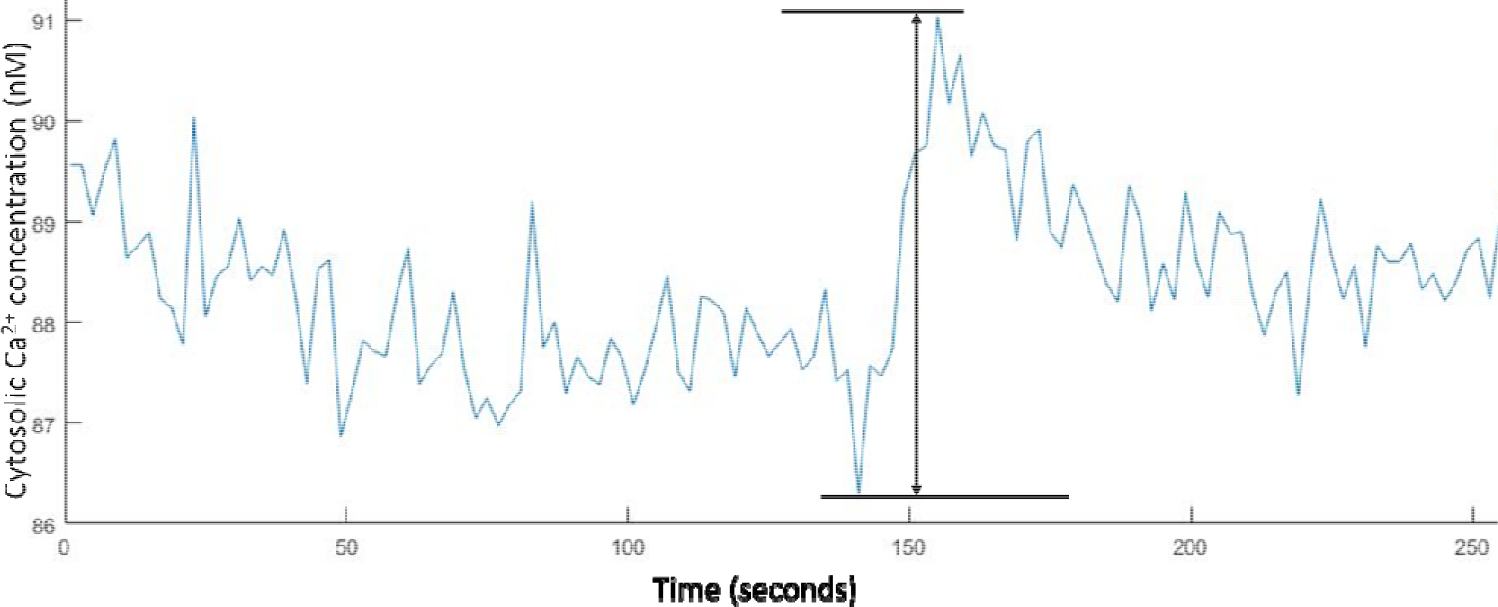
Example plot of the peak-to-peak noise recorded in the hippocampal neurons baseline Ca^2+^concentrations whilst carrying out 0.5 Hz ratiometric Fura-2 Ca2imaging.

To confirm this finding, we diluted a 17 nM free Ca^2+^ solution from a Fura-2 Ca^2+^ imaging calibration kit (Invitrogen, Paisley, UK) with distilled water to obtain a series of solutions with decreasing Ca^2+^ concentrations that decreased in steps of 2 nM. By imaging 5 µL of each solution and obtaining the background corrected fluorescence emission ratio we were able to determine that at concentrations above 5 nM we could identify changes in Ca^2+^ on the order of 2 nM.

### 340/380nm LEDs allows Ca2^2+^ imaging using lower Fura-2 AM concentrations

We investigated the possibility of using lower concentrations than the 1 µM typically recommended in Fura-2 AM loading protocols [15], [32], We imaged trypsin (100 nM) mediated Ca^2+^ transients in tsA-201 cells loaded with either 750 nM (n = 111), 500 nM (n = 119) or 250 nM (n = 130) Fura-2 AM and compared the average Ca^2+^ increase above the baseline to the value obtained in the initial experiments using 1 µM of Fura-2 AM (n = 572, Figure 6). The Ca^2+^ increases above the baseline for the 750 nM, 500 nM and 250 nM were 633.0 ± 33.9 nM (n = 111, P>0.05 compared to 1 µM Fura-2 AM) 552.6 ± 22.7 nM (n = 119, P>0.05 compared to 1 µM Fura-2 AM) and 616.0 ± 24.9 nM (n = 130, P>0.05 compared to 1 µM Fura-2 AM), respectively.

**Figure 6.**
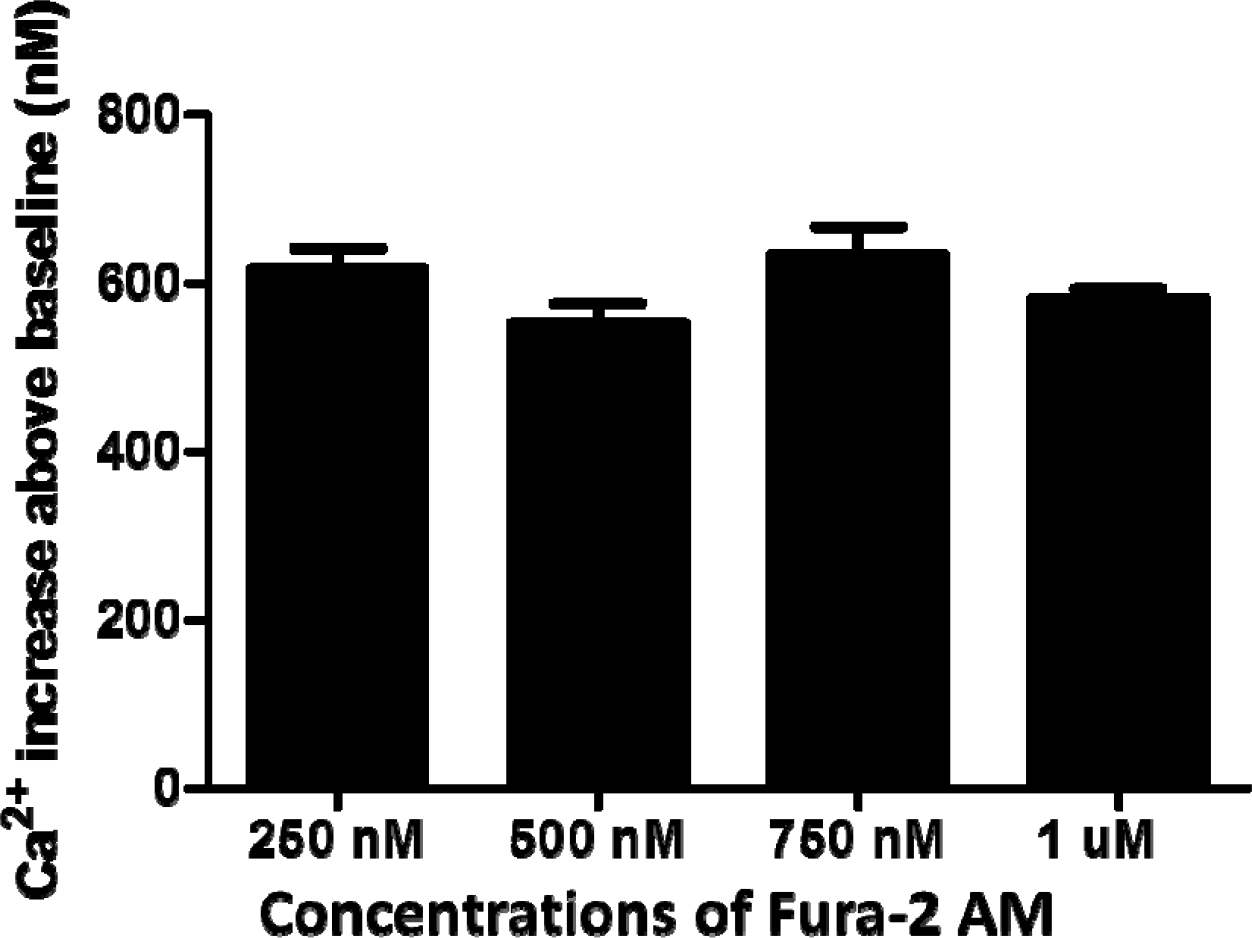
Comparison ofCa^2^’ increases obtained from the application of trypsin (100 nM) to tsA-201 cells loaded with different concentrations of Fura-2 AM.

### Video rate ratiometric Ca^2+^ imaging using 1 µM Fura-2 AM

It has been previously shown, using widefield two-photon microscopy at imaging speeds between 10 and 100 Hz, that spontaneous Ca^2+^ events could be detected in hippocampal neurons loaded with Fluo-4 AM though it was not possible to convert the fluorescence signals into a quantitative Ca^2+^ concentration [28]. Here we utilized the 340/380nm LED to image at 0.5 Hz and 24.39 Hz (limited only by the frame rate available with the camera) spontaneous synaptically driven Ca^2+^ events in hippocampal neurons, induced by the application of magnesium (Mg^2+^)-free HBS. At 0.5 Hz, a clear increase in intracellular Ca^2+^ levels were observed when Mg^2+^-free HBS was applied but individual events were difficult to decipher (Figure 7A). However, at an image acquisition rate of 24.39 Hz, individual increases in intracellular Ca^2+^ levels were observed that are similar to action potential firing seen when using patch clamping[28], with some synchronicity in firing between different neurons also being observed (Figure 7B). Indeed, a peak-to-peak measurement of the baseline fluctuations at 24.39 Hz found an average fluctuation of 7.10 ± 0.04 % (n = 21) which equates to a fluctuation in the average resting Ca^2+^ (104.5 ± 4.1 nM) of 7.42 ± 0.04 nM (n = 21). When imaging at 0.5 Hz the hippocampal neurons had an average baseline fluctuation of 5.22 ± 0.06 % (n = 39) which is a fluctuation in the basal Ca^2+^ (87.9 ± 5.3 nM) of 4.59 ± 0.05 nM.

**Figure 7.**
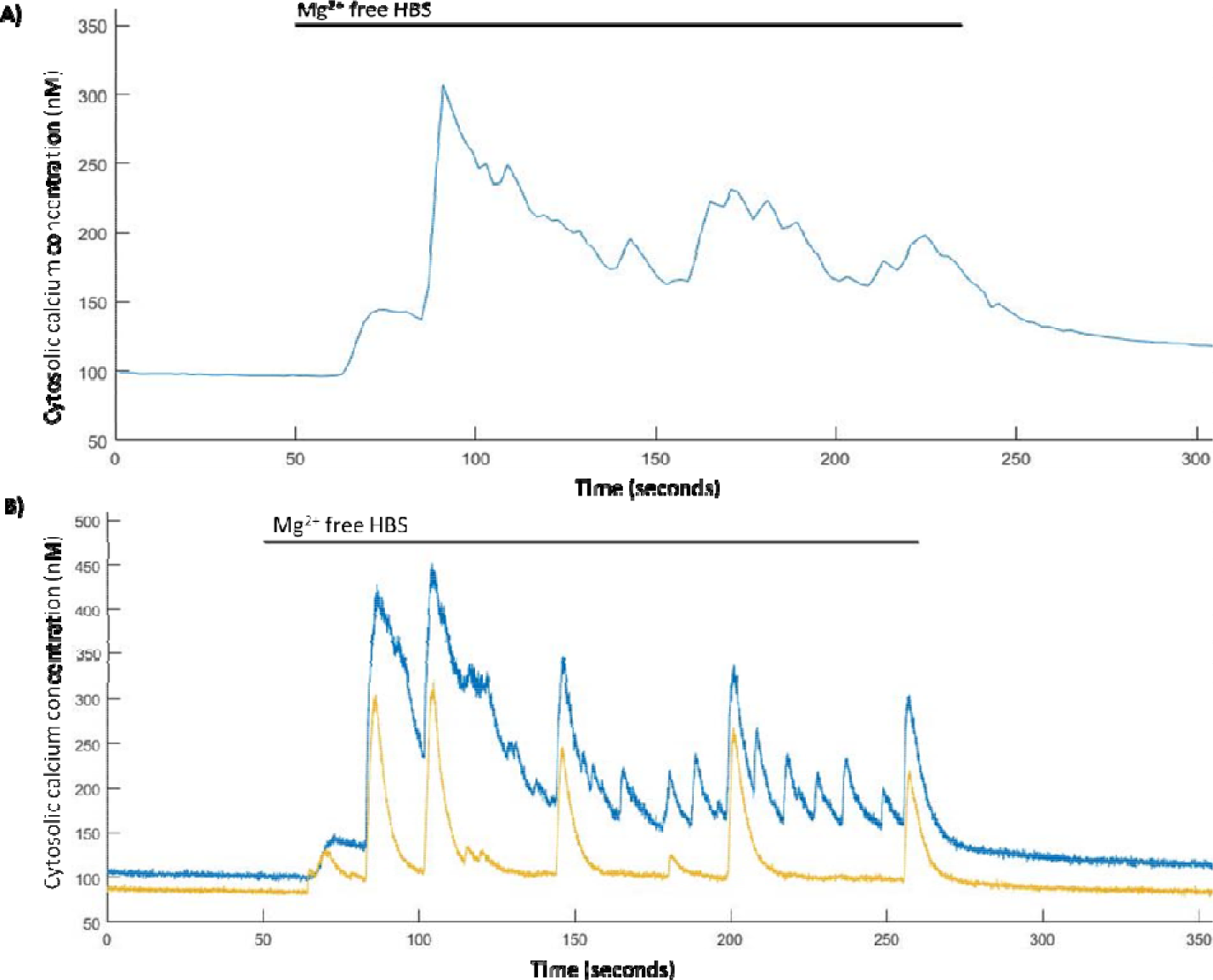
Spontaneous Ca^2^’ events are induced in Mg^2^’-free HBS. A) representative trace from a single hippocampal neuron of Mg^2^’-free induced Ca^2^’ events imaged at 0.5 Hz and B) representative trace from two hippocampal neurons of Mg^2^’-free induced Ca^2^’ events imaged at 24.39 Hz.

## Discussion

We have demonstrated the first application of a fast wavelength-switchable 340/380 nm LED illuminator for ratiometric Ca^2+^ imaging of live cells loaded with the fluorescent Ca^2+^ indicator Fura-2 AM. The 340/380 nm LEDs more accurately match the peak excitation wavelength of bound and free Ca^2+^ ions, offering more efficient excitation and higher signal-to-noise ratios when compared with other illumination systems. The fluorescence fold increases and cytosolic Ca^2+^ changes measured in the live cell specimens are in agreement with previous studies in hippocampal neurons, tsA-201 cells and HEK-293 cells illuminated using an arc lamp system[33]—[37]. Analysis of the baseline Ca^2+^ peak-to-peak noise showed that the 340/380 nm LED illuminator enables the detection of cytosolic Ca^2+^ changes with a minimum precision of 3.9 ± 0.2 nM. The limit on the precision of the experiment comes not from the imaging apparatus but only the response of the dye to Ca^2+^ which has a theoretical precision of 5 – 10 nM [38].

In addition, we have shown that using the new illumination system we were able to load cells with lower concentrations of Fura-2 AM than recommended in loading protocols with no statistically significant difference between the Fura-2 AM concentrations examined. It was possible to load our specimens with as low as 250 nM Fura-2 AM and still accurately record pharmacologically-mediated Ca^2+^ events at 0.5 Hz. The utility of lower dye concentrations presents not only an economical advantage by allowing more experiments from a single vial of dye but also may increase the viability of cells by reducing the concentrations of formaldehyde and acetic acid created through the hydrolysis of AM-ester [39].

Finally, we have demonstrated the functionality of the new illuminator system by utilizing the intrinsic LED advantages of rapid wavelength switching and amplitude stability to image and quantify synaptically-driven Ca^2+^ events in hippocampal neurons at a video rate of 24.39 Hz. We compared the Ca^2+^ traces obtained when imaging at video rate to those recorded when imaging at 0.5 Hz (Figure 7A). It can be seen from this comparison that due to the slow image capture rate of 0.5 Hz, many of the rapid synaptically driven Ca^2+^ events cannot be imaged. The appearance of these events in the presence of Mg^2+^ free HBS is due the relief of the voltage-dependent blockade of N-methyl-D-aspartate (NMDA) receptors by Mg^2+^[40], [41]. This is a well established phenomenon that is used extensively both in academia and in the pharmaceutical industry to induces synaptically driven Ca^2+^ events and eplileptiform-like activity[42]. [43]. but the clear discrimination between individual events has not previously been possible due to the limitations of arc lamp systems. It was also apparent that some synchronicity occurred between larger Ca^2+^ events in different neurons, this has been reported previously for neuronal networks under Mg^2+^ free conditions[43], [44], The increase in baseline fluctuation observed in these measurements compared to the slower imaging rates can be attributed to a decrease in the signal-to-noise ratio due to the lower exposure times used which reduces the number of photons collected for each image. Though there was an increased fluctuation it was not significantly larger from that obtained at 0.5 Hz imaging rates (P>0.05). The ability to observe these spontaneous changes in Ca^2+^ using wide-field microscopy allows for improved temporal resolution of imaging synaptically-driven neuronal Ca^2+^ events whilst maintaining the high spatial resolution afforded by Ca^2+^ imaging leading to a higher throughput of more informative measurements than currently offered with existing Ca^2+^imaging methods[6].

## Conclusions

We believe this to be the first demonstration of a truly 340/380 nm LED illuminator for ratiometric Fura-2 Ca^2+^ imaging of live cells. By matching the wavelengths of the illuminator to the optimum excitation wavelengths of the free and Ca^2+^ bound states of Fura-2, we have combined efficient ratiometric imaging with rapid switching and high intensity stability of LEDs. This represents a significant improvement over existing LED-based illuminators and frees Fura-2 ratiometric imaging from the known limitations of arc lamps.

## Acknowledgments

The authors would like to thank John Dempster for his assistance in configuring the WinFluor software for our application. Peter W. Tinning is supported by a University of Strathclyde scholarship partially funded by CoolLED Ltd. This research was also part funded by a Medical Research Council grant – MR/K015583/1.

